# Abdominal adiposity negatively associates with the rate of long term sequential skill learning

**DOI:** 10.1101/186742

**Authors:** Amira Millette, Brighid Lynch, Timothy Verstynen

**Author notes:** These authors contributed equally to this work (Amira Millette, Brighid Lynch). Corresponding Author: Timothy Verstynen, Carnegie Mellon University, Department of Psychology, 340 U Baker Hall, Pittsburgh, PA 15213, 412-533-AXN1 (office phone). **Disclosure:** The authors declare no conflict of interest.

## Abstract

Obesity is associated with functional and structural differences in the corticostriatal systems of the brain. These pathways are known to be critical for the acquisition of complex sensorimotor skills, such as the ability to learn a coordinated sequence of actions. Thus, individual differences in obesity should be associated with reduced efficiency of learning sequential skills. Here we measured long-term sequence learning across five days of training on the serial reaction time task in a cohort of neurologically healthy adults (N=30) with body types ranging from lean to obese. As expected, individuals with a greater degree of central adiposity, measured as central waist circumference, exhibited slower rates of learning, across all training days, than their leaner counterparts. This association between learning and central adiposity was restricted to response speeds, but not accuracy. These findings show that obesity is negatively associated with the efficiency of learning a long-term sequential skill, possibly due to previously observed associations between obesity and general basal ganglia function.

## 1. Introduction

Over the last ten years, a small but growing body of research suggests that obesity may be variably linked to individual differences in cognitive function. For example, higher levels of obesity have been associated with reduced efficiency in a number of cognitive domains, including inhibitory control (Duchesne et al, 2010; Galioto et al., 2012; Gunstad et al, 2007; Hendrick et al., 2012; Nederkoorn et al., 2006a; Nederkoorn et al., 2006b) and general decision making (Boeka & Lokken, 2008; Cserjési et al., 2009; Davis et al., 2004; Davis et al., 2010; Lokken et al., 2009; Pignatti et al., 2006). Indeed, the cognitive associations with obesity appear to be fairly broad. For example, in a recent study by Galioto and colleagues, participants with morbid obesity showed reduced performance in executive function, memory, language, and attention tasks, relative to lean counterparts, even in the absence of a binge eating disorder diagnosis. However, it is worth noting that while reports of obesity-cognition associations are somewhat variable, they tend to be more consistent after accounting for individual differences in mood and affect (Volkow et al., 2008a).

At the neural level, obesity is consistently associated with individual differences in the neural substrates of decision making and reward processing, specifically within a set of pathways known as the cortico-basal ganglia system (Stice et al., 2008; Tomasi & Volkow, 2013; Volkow et al., 2011). For example, individuals with high obesity have been found to have decreased levels of dopaminergic D2 receptor availability in the striatum, a set of forebrain nuclei that serve as the primary input for the basal ganglia (de Weijer et al., 2011; Volkow et al., 2008b). Functional imaging studies also show that, compared to their lean counterparts, individuals with high adiposity have a hypo-responsive ventral striatal reward response that is thought to lead to overeating as a compensation for this blunted dopaminergic signal (Stice et al., 2008; Stice et al., 2010). These associations with striatal function are often discussed in the context of reward and hedonic overeating (Horstmann et al., 2011); however, the cortico-basal ganglia pathways also play critical roles in executive control and learning, suggesting that obesity may also be associated with non-reward related cognitive functions.

One cognitive function that relies critically on the integrity of cortico-basal ganglia pathways is sequential skill learning (Doyon et al., 2009; Lehericy et al., 2005). This is the cognitive ability to bind temporally independent items together into a unified sequence and is critical to many everyday behaviors, e.g., driving a vehicle with a manual transmission, learning a language, playing a new piano melody. Neuroimaging studies have shown that regions within the basal ganglia, particularly the striatal nuclei, are engaged during sequence learning and their activity is modulated over the course of training (Lehericy et al., 2005). Causal evidence for the role of striatal systems in motor sequence learning comes from studies of neurological patient populations with pathologies of the basal ganglia. For example, Parkinson’s disease is a degenerative disorder of the dopaminergic projections from the substantia nigra pars compacta to the striatal nuclei. Parkinson’s disease patients show significant deficits in motor sequence skill acquisition and consolidation when introduced to a novel motor sequence (Benecke et al., 1987; Dan et al., 2015). This patient evidence, combined with the neuroimaging evidence, highlights the critical role that striatal nuclei play in sequence acquisition (Doyon, 2008).

Since obesity is associated with reduced striatal functioning and the striatum is critical to motor sequence learning, then it is possible that obesity is associated with the efficiency of learning a novel sequential sensorimotor skill; however, this association has yet to be tested. The current project aims to fill this gap by using a standard inventory of motor sequence learning, the serial reaction time task (Nissen & Bullemer, 1987), to measure the rate of long-term learning across five days of training in a sample of neurologically healthy participants with body types ranging from lean to obese. Specifically, we predict that individual differences in central adiposity, measured as waist circumference, would negatively correlate with the ability to learn a complex sequence of actions across multiple days of training.

## 2. Methods and Materials

### 2.1 Participants

Thirty-two right-handed volunteers were recruited from the local Pittsburgh community with a body-mass index range of 18.5kg/m^2^ to 40.0kg/m^2^. While relatively small for biomedical and health psychology studies of obesity, our sample size is quite consistent with typical SRTT studies on both clinical and healthy populations (for example see Knopman & Nissen, 1991; Shin & Ivry, 2003). All participants reported no more than three years of musical experience in the last 10 years. Additional inclusion criteria included that participants reported current, unimpeded use of the right hand, had no history of carpal tunnel syndrome or similar disorders, and no familiarity with the Cyrillic alphabet. Each volunteer gave written, informed consent, and were financially compensated for their participation. Two participants failed to attend all five consecutive days of training and were therefore removed from the final data set. All procedures were approved by the Carnegie Mellon University Institutional Review Board (IRB) prior to testing.

### 2.2 Obesity measures

Body mass index was used as the primary measure of obesity for recruitment purposes. Height and weight measurements were taken from each participant prior to the start of the experimental task. Body mass index was calculated using the standard formula of weight in kilograms (kg) divided by the square of height in meters (m^2^). The standard ranges of body mass index were categorized as, lean (BMI of 18.5 - 24.9), overweight (BMI of 25.0 - 29.9), and obese (BMI greater than 30.0).

Waist circumference was used as a direct measure of central adiposity in the statistical analysis. The waist circumference measurement was taken from each participant prior to the start of the experimental task, using a plastic tape to measure around the participant’s waist, just above the navel. Since there is no established definition of obesity based on waist circumference, we performed a median split on the waist circumference measure to create a categorical variable as the new waist circumference measure, with 36.0 inches as the median value. The lowest waist circumference value to 36.0 inches served as the “Low Adiposity” group (N=16), and 36.1 inches to the highest waist circumference value in the dataset served as the “High Adiposity” group (N=14). This categorization did not align perfectly with BMI categories, with 9 lean, 6 overweight, and 1 obese individuals in the Low Adiposity group and 1 lean, 4 overweight, and 9 obese individuals in the High Adiposity group.

### 2.3 Serial Reaction Time Task

#### 2.3.1. Experimental Design and Setup

Participants were run in a standard version of the serial reaction time task for five consecutive days, with a one-hour training session on each day. All stimuli were presented on a 23” ASUS LED monitor with a resolution of 1920 x 1080 mp, using Matlab R2012a (MathWorks, Inc., Natick, MA). The stimuli were spatially centered on the computer screen in a black background, displayed in a white font color (Fig. 1). Participants were told to respond to a set of cued stimuli presented visually on the computer screen using the right index (1), middle (2), ring (3), and pinky (4) fingers consecutively, with each finger matched to one uniquely paired cue in this order: “ж”, “Є”, “Њ”, and “Л” (Fig. 1A). Each experimental session consisted of a total of six trial blocks. The experimental blocks were divided into types: Random blocks and Sequence blocks (see Fig. 2A-B). The Random blocks (trial blocks 1, 2, and 5) were comprised of 264 stimuli presented in a pseudorandom order, with a restriction to minimize repetition between any two contiguous stimuli. The Sequence blocks (trial blocks 3, 4, and 6) were comprised of twenty-two repetitions of stimuli from a 12-trial sequence [1, 3, 2, 1, 4, 3, 1, 4, 2, 3, 4, 2]. Each block began at a random part of the sequence so as to minimize immediate identification of the sequential pattern. The sequence pattern remained the same for all Sequence blocks and across the five training days. The experiment was self-paced such that participants were allowed to proceed to the next trial block when they were prepared for the following series.

**Figure 1:**
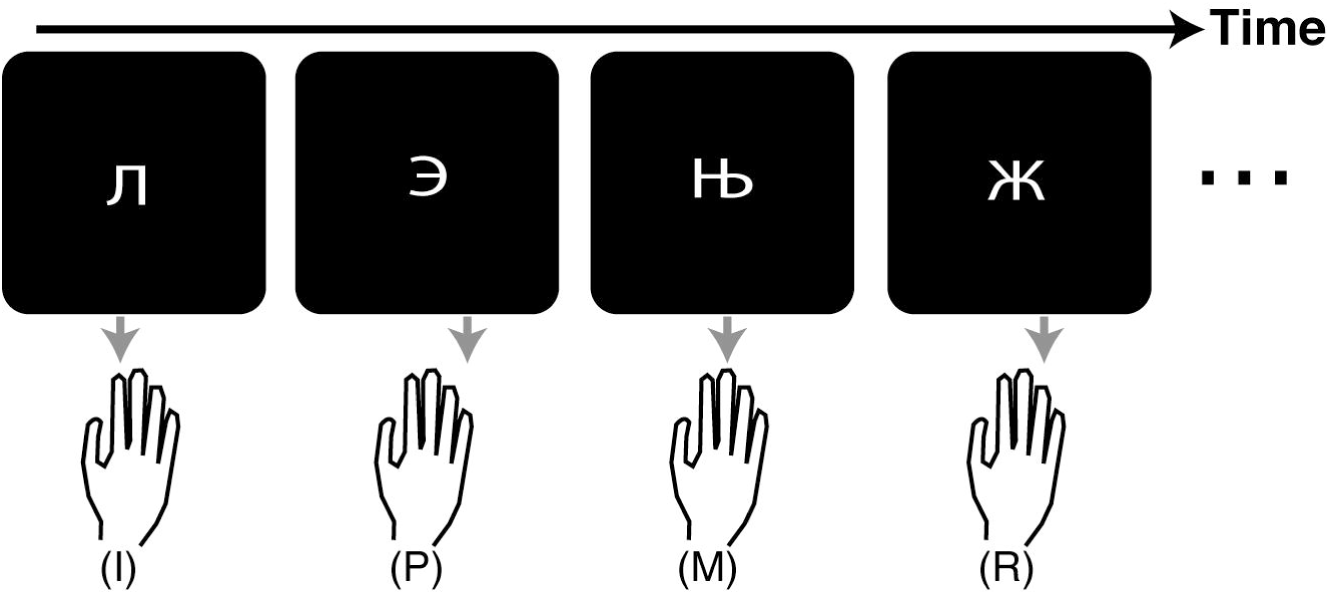
Block-wise mean a) response times (RT) and b) accuracies across experimental cohort. All error bars are standard error of the mean.

**Figure 2:**
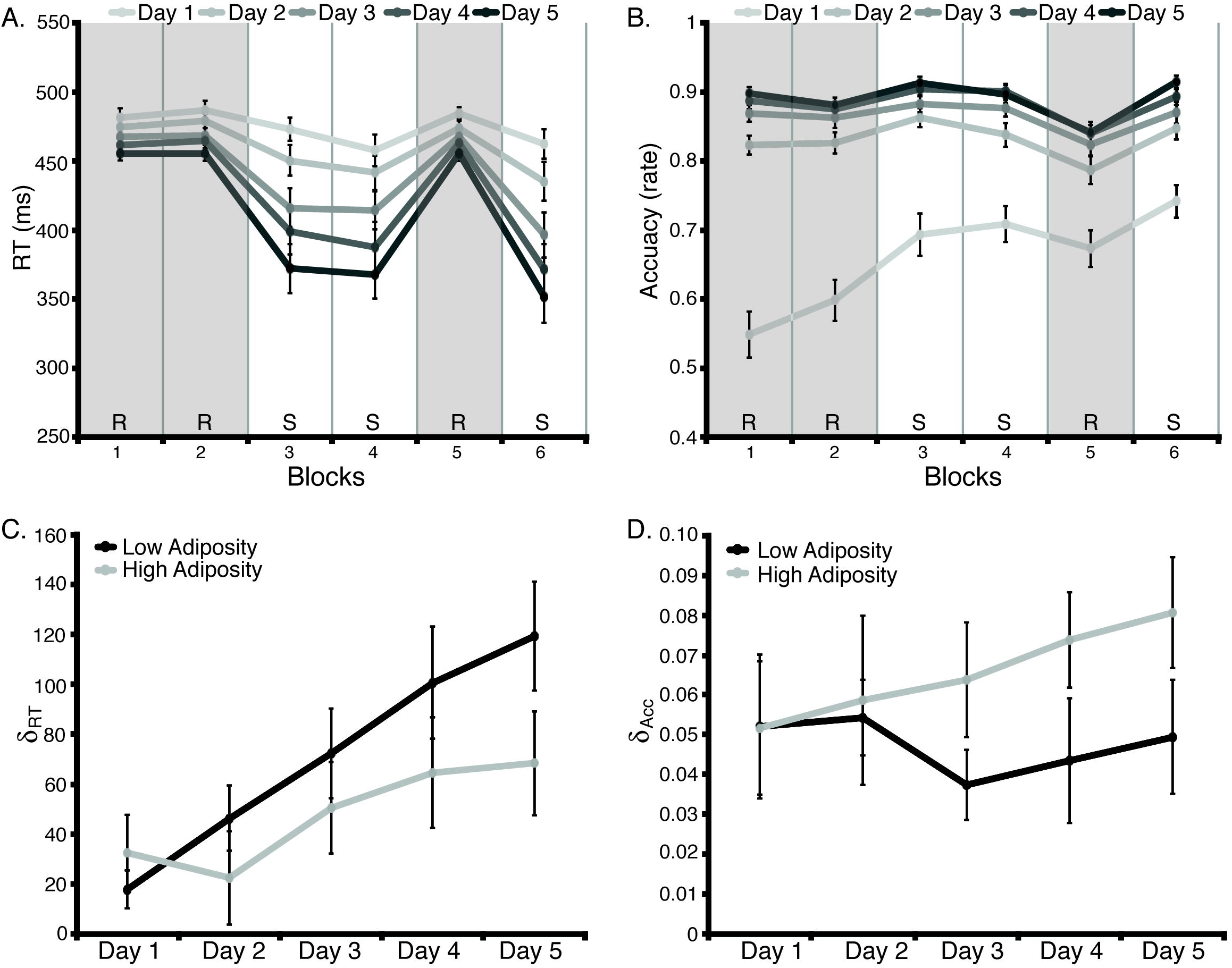
Learning across training days. Mean blockwise scores for a) response time and b) accuracies across all subjects and days of training. Sequence specific learning scores (δ) on each day of training (see main text for details) for c) response time and d) accuracy, separated by adiposity group. All error bars are standard error of the mean.

#### 2.3.2 Task Instructions

Prior to the start of the experimental session, participants were informed that there was a 600ms response window and they were instructed to respond as quickly and as accurately as possible. Participants then were given a brief 12 trial practice session to familiarize themselves with the key and cue mappings. After this brief practice, the testing session began. Throughout the training session, participants were provided continuous visual feedback on their response accuracy; a correct key press resulted in the cue flashing green. Alternatively, an incorrect key response resulted in the cue flashing red. At the end of each trial block, participants were provided a feedback summary on their average response times and accuracies as a percent correct score for that block.

### 2.4 Data analysis

#### 2.4.1. Analysis measures

The raw data was summarized for each block using a custom Matlab script. Mean response times (response time; in ms) and accuracies (% correct) were measured on each block to determine motor sequence learning in the task. These measures were used to compute the learning scores of each body type category, across the five experimental sessions.

#### 2.4.2 Learning scores

Sequence specific learning was measured for response time by taking the mean value in the last two sequence probes (Block 4 and Block 6; μ^4^, μ^6^) and subtracting those from the same values from the last random probe (Block 5; μ^5^). This measure provided information on sequence specific learning. Learning scores in response time were manually computed (eq. 1) by subtracting the mean response time in the random probe from the average of the mean response times of the sequence probes:

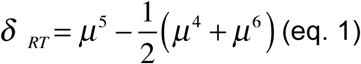

Likewise, learning scores in accuracy were manually computed (eq. 2) by taking the average of the accuracy of the random probes (α^4^, α^6^) and subtracting the accuracy in the sequence probe (α^5^):

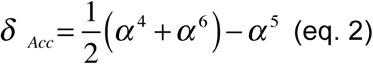

Group differences in δ_RT_ and δ_Acc_ were determined using a two-way repeated measures ANOVA.

#### 2.4.3. Learning slope

Finally, in order to quantify the linear rate of learning across training days, a learning slope was calculated (eq. 3) for each subject on sequence specific differences in response times and accuracies separately. We computed this by calculating the average between day difference in learning scores, for both response times and accuracies. This was computed as

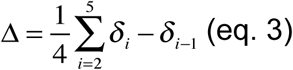

where δ_i_ is the learning score on day *i*. This score reflects the mean change in learning, per day across the entire experiment. This value was correlated against anthropometric measures of obesity using a Spearman’s rank order correlation coefficient.

## 3. Results

### 3.1. Response Time and Accuracy

Mean response times and accuracies were recorded for each block of trials for all days. Figure 2A shows the block-wise response times averaged across all subjects for the five training days. There is a noticeable increase in response speeds during the sequence probes (Blocks 4 and 6), compared to the random probe (Block 5). Across most blocks there is also a saturation of the accuracy after Training Day 1 and this was particularly strong in the sequence blocks (Fig. 2B). The last random block provides a “learning probe” that allows for us to measure sequence-specific learning at the end of each day. We calculated learning scores based off of these probe blocks for each subject and each day (see Methods).

### 3.2. Learning Score

In order to look for group differences in learning scores for response time and accuracy, we performed a median split on our main obesity variable of interest, waist circumference, and categorized subjects into Low (N=16) and High (N=14) adiposity groups. A repeated-measures ANOVA found a significant main effect of training day on response time learning scores (Fig. 2C; *F* (4, 116) = 17.55; *p* < 0.001), as well as a significant group by day interaction (*F* (4, 116 = 2.51; *p* = 0.045). In general, the Low Adiposity group had a greater rate of learning across training than the High Adiposity group. This group effect is driven by across day learning, as 2-sample t-tests did not find a significant group difference on each individual training day (all p’s > 0.10, all t’s < 1.68). This association between sequence learning and abdominal adiposity group appears to be specific for movement speeds. The learning scores based on accuracy performance did not show a training day by group interaction (Fig. 2D; *F* (4, 116) < 1; *p* = 0.565). It is noteworthy that this effect of body type on long-term learning is not significant when BMI group is used as the group category as there was not a significant interaction between response time learning score and body-mass index (*F* (2, 116) = 0.327; *p* =0.540). This is consistent with previous results showing that waist circumference is as a more reliable and direct measure of obesity than BMI (Gómez-Ambrosi, et al. 2012).

### 3.3. Learning Slope

To better understand the relationship between central adiposity and response time learning, we used a linear slope analysis to estimate the rate of across day learning for each subject (see Methods). Using a non-parametric Spearman’s rank correlation test (*ρ*_S_), we found a significant negative correlation between learning slope of response times and waist circumference (Table 2; *ρ*_S_ = −0.401, *p* < 0.05). Consistent with our hypothesis, as central adiposity increased, the rate of motor sequence learning across days decreased (Fig. 3). Within each day, we saw the emergence of an association between central adiposity and sequence specific response time learning, becoming significant on the last day of training (Day 1: *ρ*_S_ = 0.13, *p* = 0.40; Day 2: *ρ*_S_ = −0.14, *p* = 0.07; Day 3: *ρ*_S_ = −0.04, *p* = 0.23; Day 4: *ρ*_S_ = −0.12, *p* = 0.09; Day 5: *ρ*_S_ = −0.25, *p* = 0.03). There was a similarly negative correlation between learning slope of accuracies and BMI but it did not reach statistical significance (*ρ*_S_ = −0.230, *p* = 0.221). As with response time learning, an association between central adiposity and sequence specific accuracy learning emerged on each day, with a significant association on the last day of training (Day 1: *ρ*_S_ = 0.21, *p* = 0.22; Day 2: *ρ*_S_ = −0.03, *p* = 0.35; Day 3: *ρ*_S_ = 0.29, *p* = 0.10; Day 4: *ρ*_S_ = 0.35, *p* = 0.05; Day 5: *ρ*_S_ = 0.38, *p* = 0.04).

**Table 2:** Spearman’s non-parametric correlation coefficient between each variable and covariate. Body mass index, BMI; Waist circumference; WC; Learning score, δ; Response time, RT; Accuracy, Acc.

| | Sex | Age | BMI | WC | $\delta_{RT}$ Slope |
| --- | --- | --- | --- | --- | --- |
| Age | -<br>0.035 | - | - | - | - |
| BMI | -<br>0.054 | 0.257 | - | - | - |
| WC | -<br>0.327 | 0.357 | 0.740** | - | - |
| $\delta_{RT}$ Slope | .0307 | -<br>0.236 | -0.230 | -0.401* | - |
| $\delta_{Acc}$ Slope | -<br>0.023 | -<br>0.178 | 0.103 | 0.191 | -0.115 |

\* Significance at $\alpha < 0.01$
\*\* Significance at $\alpha < 0.05$

**Figure 3:**
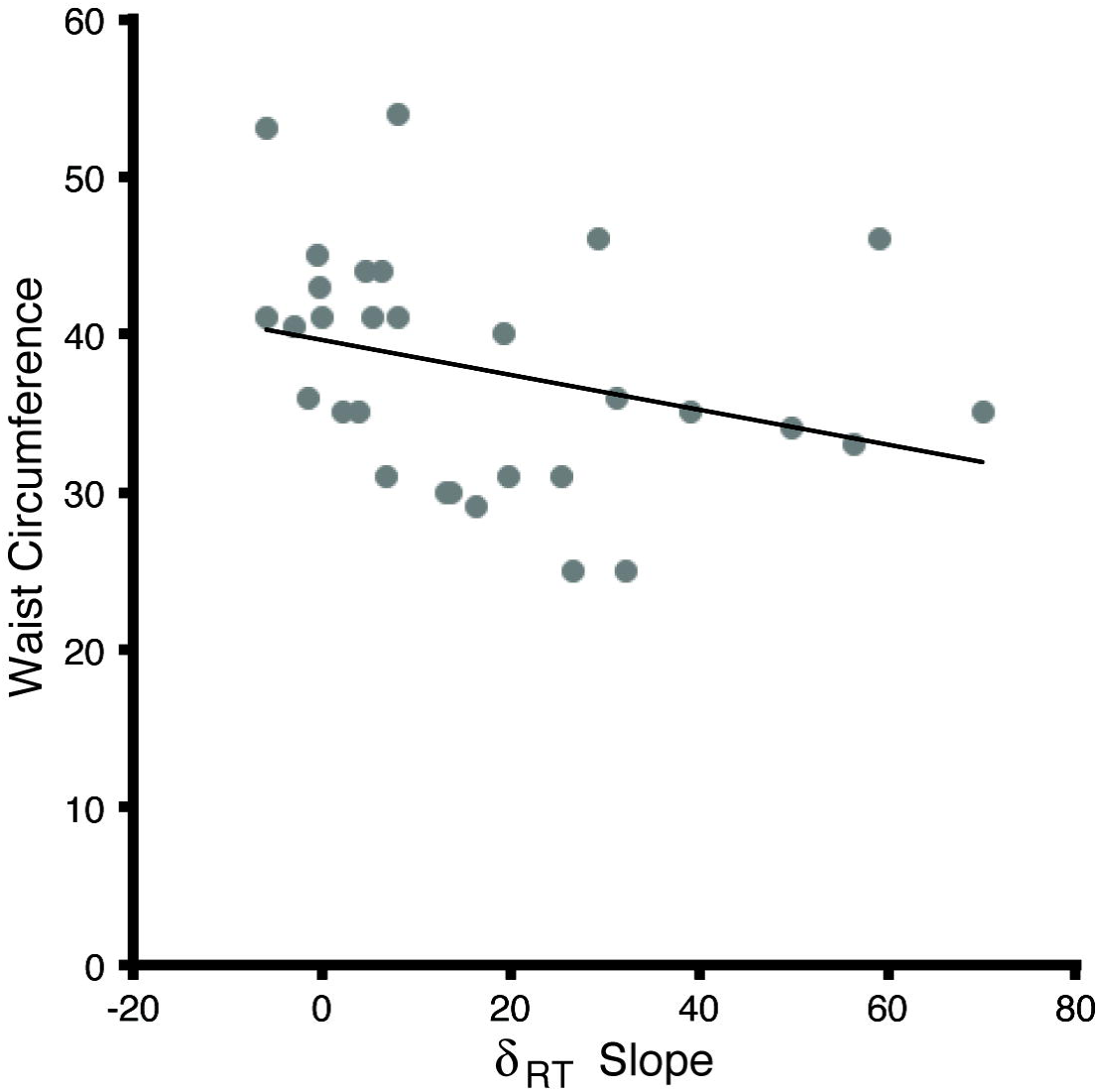
Individual differences in learning score slope for response time (δ_RT_ Slope) and waist circumference.

## 4. Discussion

For the first time we have shown that a measure of obesity, central adiposity, is associated with a decrease in the efficiency of long-term sequential skill learning. Participants with a higher waist circumference acquired a sensorimotor sequence at a slower rate than lean counterparts across five days of training on the serial reaction time task. This association with central adiposity was limited to response speeds, but not accuracy, consistent with the idea that these two performance measures may be learned independently (see also Verstynen et al., 2012).

Given the critical role of basal ganglia networks in long term skill learning (Doyon, 2008), these findings fit with the growing body of research that obesity is related to the integrity and efficiency of basal ganglia circuits. Much of this analysis has focused on the relationship between obesity and reactivity of striatal reward pathways (Tomasi & Volkow, 2013; Wang et al., 2001). For example, Stice and colleagues found a negative association between dorsal striatal reactivity in response to a high value food stimulus and BMI. Importantly, this was moderated by the presence of the dopamine allele of the TaqIA polymorphism. In addition, individuals with high obesity also exhibit reduced dopamine D2 receptor availability throughout the striatum when compared to the lean counterparts (de Weijer et al., 2011; Volkow et al., 2008). Our finding that obesity is associated with reduced efficiency of a striatal-dependent form of learning, suggests these striatal associations with obesity may relate to global functioning of basal ganglia pathways, rather than just processing the saliency of rewarding stimuli. Specifically, reward prediction errors are coded by phasic dopaminergic signals into the striatum and these reward prediction errors are crucial for reinforcement learning (Frank & Badre, 2012). Blunted dopamine reactivity in the striatum of individuals with obesity would reduce the sensitivity of this learning signal over time, thereby slowing the rate of learning. Thus, individual differences in dopamine may be the primary mechanism impacting the rate of skill learning reported here. Future work should focus on neural measures of reward saliency and how they relate to learning within the context of obesity.

Our findings, considered in the context of previous striatum-obesity associations (Horstmann et al., 2010; Nummernmaa et al., 2012; Stice et al., 2008; Tomasi & Volkow, 2013), suggests that obesity may reflect an inherent concern of basal ganglia function. Although there is some heterogeneity in reports of obesity-cognition links (Fitzpatrick et al., 2013), the most reliable cognitive associations with obesity tend to be those processes (e.g., inhibitory control, reward saliency, decision making) that also rely on healthy cortico-basal ganglia pathway function. This observation highlights a system, cortico-basal ganglia networks, that could better explain the specific behavioral patterns that lead to increased obesity.

While our results suggest that obesity itself may associate with long-term procedural learning, many other factors correlate with obesity that may also contribute to the rate of motor learning. For example, obesity is correlated with reduced cardiorespiratory fitness (Ross & Katznarzyk, 2003; Wong et al., 2004). Accumulating evidence suggests that cardiorespiratory fitness impacts cognitive function (Szabo et al., 2011). Since we did not measure cardiorespiratory fitness in this study, we cannot preclude this possibility here. Beyond cardiorespiratory fitness, many physiological (e.g., inflammation) and metabolic (e.g., insulin) systems associated with elevated adiposity have also been associated with variability in cognitive function (Rosano et al., 2012). Understanding how these physiological and metabolic factors may moderate or mediate the association between adiposity and skill learning should be a focus of future studies. Testing this requires replicating the current study with a much larger sample size that includes additional measurements of health related systems.

Of course, the current association between central adiposity and skill learning could be explained by secondary factors not related to basal ganglia function per se. For example, participants with higher central adiposity may also fatigue sooner than their lean counterparts, thus arousal levels may also account for efficiency of motor learning. Follow-up work should quantify levels of fatigue across groups as a control for this possibility. Finally, obesity has been shown to be comorbid with depression and anxiety disorders, which may impact attention and, consequently, learning (Strine et al., 2008). Administering a standardized battery to quantify mood and affect will aid in controlling for these additional extrinsic factors. Follow-up work should also confirm the that the striatum is mediating the relationship between obesity and long-term motor sequence skill learning by looking at striatal activity during this form of long-term learning across groups. Neuroimaging tools, such as fMRI, can provide valuable information to the particular regions involved in this motor learning task, and with this, we can confirm the striatal link to obesity to motor sequence skill learning.

Regardless of these mechanistic limitations, our results show, for the first time, that individual differences in obesity levels negatively associate with long-term skill learning. This extends our understanding of obesity and cognition links beyond reward processing and implies that obesity may associate with general basal ganglia function. This has wide-ranging implications on the role of obesity in complex cognitive functions, like sensorimotor skill learning, and opens new avenues of research into the effects of physical fitness on the brain beyond reward processing.

**Table 1:** Participant demographics.

|  | Lean (n = 10) | Overweight (n = 10) | Obese (n = 10) |
| --- | --- | --- | --- |
| Sex |  |  |  |
| Male | 4 | 8 | 5 |
| Female | 6 | 2 | 5 |
| Age (in years) |  |  |  |
| Mean | 21.1 | 22.8 | 25.4 |
| Median | 20.5 | 21.5 | 23.5 |
| St. Dev. | 1.91 | 5.37 | 8.25 |
| Minimum | 19 | 18 | 19 |
| Maximum | 25 | 36 | 46 |
| BMI (kg/m<sup>2</sup>) |  |  |  |
| Mean | 21.91 | 27.22 | 32.52 |
| Median | 22.9 | 27.4 | 31.55 |
| St. Dev. | 2.23 | 1.53 | 2.66 |
| Minimum | 18.1 | 25.0 | 30.4 |
| Maximum | 24.4 | 29.0 | 34.1 |
| Waist Circumference<br>(in inches) |  |  |  |
| Mean | 31.1 | 38.3 | 43.7 |
| Median | 30.5 | 35.5 | 44.0 |
| St. Dev. | 4.79 | 6.46 | 4.67 |
| Minimum | 25.0 | 31.0 | 35.0 |
| Maximum | 41.0 | 54.0 | 53.0 |

## Acknowledgments

This research was funded in part by a grant from the Pennsylvania Department of Health’s Commonwealth Universal Research Enhancement Program #SAP4100062201 and a National Science Foundation CAREER Award #1351748.

## References

Benecke R, Rothwell JC, Dick JPR, Day BL, Marsden CD. Disturbance of sequential movements in patients with Parkinson’s disease. Brain 1987;110:361–379.

Boeka AG, Lokken KL. Neuropsychological performance of a clinical sample of extremely obese individuals. Arch Clin Neuropsychol 2008;23:467–474.

Cserjési R, Luminet O, Poncelet AS, Lénárd L. Altered executive function in obesity. Exploration of the role of affective states on cognitive abilities. Appetite 2009;52:535–539.

Dan X, King BR, Doyon J, Chan P. Motor sequence learning and consolidation in unilateral de novo patients with Parkinson’s disease. PLoS One 2015;10:1–14.

Davis C, Levitan RD, Muglia P, Bewell C, Kennedy JL. Decision-making deficits and overeating: a risk model for obesity. Obes Res 2004;12:929–935.

Davis C, Patte K, Curtis C, Reid C. Immediate pleasures and future consequences. A neuropsychological study of binge eating and obesity. Appetite 2010;54:208–213.

de Weijer B a, van de Giessen E, van Amelsvoort T a, et al. Lower striatal dopamine D2/3 receptor availability in obese compared with non-obese subjects. EJNMMI Res 2011;1:37.

Doyon J. Motor sequence learning and movement disorders. Curr Opin Neurol 2008;21:478–83.

Doyon J, Bellec P, Amsel R, et al. Contributions of the basal ganglia and functionally related brain structures to motor learning. Behav Brain Res 2009;199:61–75.

Duchesne M, Mattos P, Appolinário JC, et al. Assessment of executive functions in obese individuals with binge eating disorder. Rev Bras Psiquiatr 2010;32:381–388.

Fitzpatrick S, Gilbert S, Serpell L. Systematic review: Are overweight and obese individuals impaired on behavioural tasks of executive functioning? Neuropsychol Rev 2013;23:138– 156.

Frank MJ, Badre D. Mechanisms of hierarchical reinforcement learning in corticostriatal circuits 1: Computational analysis. Cereb Cortex 2012;22:509–526.

Galioto R, Spitznagel MB, Strain G, et al. Cognitive function in morbidly obese individuals with and without binge eating disorder. Compr Psychiatry 2012;53:490–495.

Gómez-Ambrosi J, Silva C, Galofré JC, et al. Body mass index classification misses subjects with increased cardiometabolic risk factors related to elevated adiposity. Int J Obes 2012;36:286–294.

Gunstad J, Paul RH, Cohen RA, Tate DF, Spitznagel MB, Gordon E. Elevated body mass index is associated with executive dysfunction in otherwise healthy adults. Compr Psychiatry 2007;48:57–61.

Hendrick OM, Luo X, Zhang S, Li CR. Saliency processing and obesity: a preliminary imaging study of the stop signal task. Obesity 2012;20:1796–802.

Horstmann A, Busse FP, Mathar D, et al. Obesity-Related Differences between Women and Men in Brain Structure and Goal-Directed Behavior. Front Hum Neurosci 2011;5:58.

Knopman D, Nissen MJ. Procedural learning is impaired in Huntington’s disease: Evidence from the serial reaction time task. Neuropsychologia 1991;29:245–254.

Lehericy S, Benali H, De MP Van. Distinct basal ganglia territories are engaged in early. PNAS 2005;102.

Lokken KL, Boeka AG, Austin HM, Gunstad J, Harmon CM. Evidence of executive dysfunction in extremely obese adolescents: a pilot study. Surg Obes Relat Dis 2009;5:547–552

Nederkoorn C, Braet C, Van Eijs Y, Tanghe A, Jansen A. Why obese children cannot resist food: The role of impulsivity. Eat Behav 2006a;7:315–322.

Nederkoorn C, Smulders FTY, Havermans RC, Roefs A, Jansen A. Impulsivity in obese women. Appetite 2006b;47:253–256.

Nissen MJ, Bullemer P. Attentional requirements of learning: Evidence from performance measures. Cogn Psychol 1987;19:1–32.

Nummenmaa L, Hirvonen J, Hannukainen JC, et al. Dorsal striatum and its limbic connectivity mediate abnormal anticipatory reward processing in obesity. PLoS One 2012;7:e31089.

Pignatti R, Bertella L, Albani G, Mauro A. Decision-making in obesity: A study using the gambling task. Eat Weight Disord Anorexia, Bulim Obes 2006;11:126–132.

Rosano C, Marsland AL, Gianaros PJ. Maintaining brain health by monitoring inflammatory processes: a mechanism to promote successful aging. Aging Dis 2012;3:16–33.

Ross R, Katzmarzyk PT. Cardiorespiratory fitness is associated with diminished total and abdominal obesity independent of body mass index. Int J Obes Relat Metab Disord 2003;27:204–210.

Shin JC, Ivry RB. Spatial and temporal sequence learning in patients with Parkinson’s disease or cerebellar lesions. J Cogn Neurosci 2003; 15:1232–1243. Strine TW, Mokdad AH, Dube SR, et al. The association of depression and anxiety with obesity and unhealthy behaviors among community-dwelling US adults. Gen Hosp Psychiatry 2008; 30: 127-137.

Stice E, Spoor S, Bohon C, Small DM. Relation between obesity and blunted striatal response to food is moderated by TaqIA A1 allele. Science (80-) 2008;322:449–52.

Stice E, Yokum S, Blum K, Bohon C. Weight gain is associated with reduced striatal response to palatable food. J Neurosci 2010;30:13105–9.

Szabo AN, McAuley E, Erickson KI, et al. Cardiorespiratory fitness, hippocampal volume, and frequency of forgetting in older adults. Neuropsychology 2011;25:545–53.

Tomasi D, Volkow ND. Striatocortical pathway dysfunction in addiction and obesity: differences and similarities. Crit Rev Biochem Mol Biol 2013;48:1–19.

Verstynen TD, Phillips J, Braun E, Workman B, Schunn C, Schneider W. Dynamic Sensorimotor Planning during Long-Term Sequence Learning: The Role of Variability, Response Chunking and Planning Errors. PLoS One 2012;7:e47336.

Volkow ND, Wang G-J, Fowler JS, Telang F. Overlapping neuronal circuits in addiction and obesity: evidence of systems pathology. Philos Trans R Soc Lond B Biol Sci 2008a;363:3191–3200.

Volkow ND, Wang G-J, Telang F, et al. Low dopamine striatal D2 receptors are associated with prefrontal metabolism in obese subjects: possible contributing factors. Neuroimage 2008b;42:1537–43.

Volkow ND, Wang GJ, Fowler JS, Tomasi D, Baler R. Food and Drug Reward: Overlapping Circuits in Human Obesity and Addiction. Curr Top Behav Neurosci 2011. Wang GJ, Volkow ND, Logan J, et al. Brain dopamine and obesity. Lancet 2001;357:354–357.

Wong SL, Katzmarzyk P, Nichaman MZ, Church TS, Blair SN, Ross R. Cardiorespiratory fitness is associated with lower abdominal fat independent of body mass index. 2004.

